# Label-free quantitative phenotyping of hepatic stellate cell activation using holotomography with AI-enabled subcellular segmentation

**DOI:** 10.64898/2026.01.26.701682

**Authors:** Sin-hyoung Hong, Junhyung Park, Haesoo Kim, Hyunyoong Moon, Keehang Lee, Hana Lee, Sumin Lee, YongKeun Park

## Abstract

Hepatic stellate cell (HSC) activation is a central driver of liver fibrosis, yet its quantitative characterization in living cells remains limited by endpoint assays that rely on fixation and labeling. Here we introduce a label-free framework that combines three-dimensional holotomography (HT) with automated, AI-assisted analysis to study HSC activation dynamics in live cells. Using refractive index tomography, we non-invasively visualize hallmark structural features of HSC activation, including lipid droplet depletion, cytoskeletal remodeling, and changes in cellular morphology. Correlative fluorescence imaging validates the biological relevance of HT-derived features, while longitudinal imaging reveals continuous activation trajectories at single-cell resolution. Automated segmentation of whole cells and subcellular organelles enables scalable extraction of multi-parametric biophysical descriptors, defining a quantitative phenotypic fingerprint that distinguishes quiescent and activated states. Together, this work establishes holotomography-based quantitative phenotyping as a powerful approach for studying HSC activation and, more broadly, dynamic cell-state transitions in living systems.

## 1. Introduction

Quantitative characterization of dynamic cell-state transitions remains a central challenge in modern cell biology [1–3]. Many biologically and clinically relevant processes— including differentiation, activation, and drug response— unfold gradually over time and involve coordinated changes in cellular morphology, intracellular organization, and metabolic state. However, most widely used experimental approaches rely on endpoint assays that provide only static snapshots of these processes, limiting access to temporal trajectories and introducing perturbations through fixation or labeling [4].

Label-free imaging techniques offer a promising alternative by enabling direct observation of intrinsic cellular properties without exogenous markers. Among these, quantitative phase imaging (QPI) has emerged as a powerful approach for measuring optical path length variations that reflect cellular mass density and organization [5–7]. Recent advances in three-dimensional implementations of QPI, collectively referred to as holotomography (HT), have substantially expanded the accessible information space by enabling volumetric reconstruction of refractive index (RI) distributions in living cells [8–13]. HT has been exploited in cell biology [14–30], 3D biology [24,31–45], cancer biology [17,22,46–63], and microbiology [64–72]. These developments have transformed HT from a contrast-enhanced imaging modality into a quantitative platform capable of probing cellular structure and composition in three dimensions.

Despite these advances, two key limitations have constrained the broader adoption of label-free imaging for phenotypic analysis. First, the lack of systematic validation against established molecular markers has raised questions about the biological interpretability of RI-based features. Second, extracting quantitative, scalable phenotypic descriptors from complex three-dimensional datasets has traditionally required manual analysis or task-specific heuristics, limiting throughput and reproducibility. Addressing these challenges is essential for extending label-free imaging from qualitative observation to standardized phenotypic profiling [73].

Hepatic stellate cell (HSC) activation provides a compelling model system to address these challenges [74–76] (Fig. 1A). In healthy liver tissue, quiescent HSCs function as vitamin A–storing cells characterized by abundant lipid droplets and compact morphology. Upon chronic injury or profibrotic stimulation, HSCs undergo activation into myofibroblast-like cells, accompanied by lipid droplet depletion, cytoskeletal remodeling, and increased extracellular matrix production [77]. Although these hallmarks are well established, their progression is typically assessed using fixed-cell immunostaining at discrete time points, obscuring activation dynamics and limiting quantitative comparison across conditions [78].

**Figure 1.**
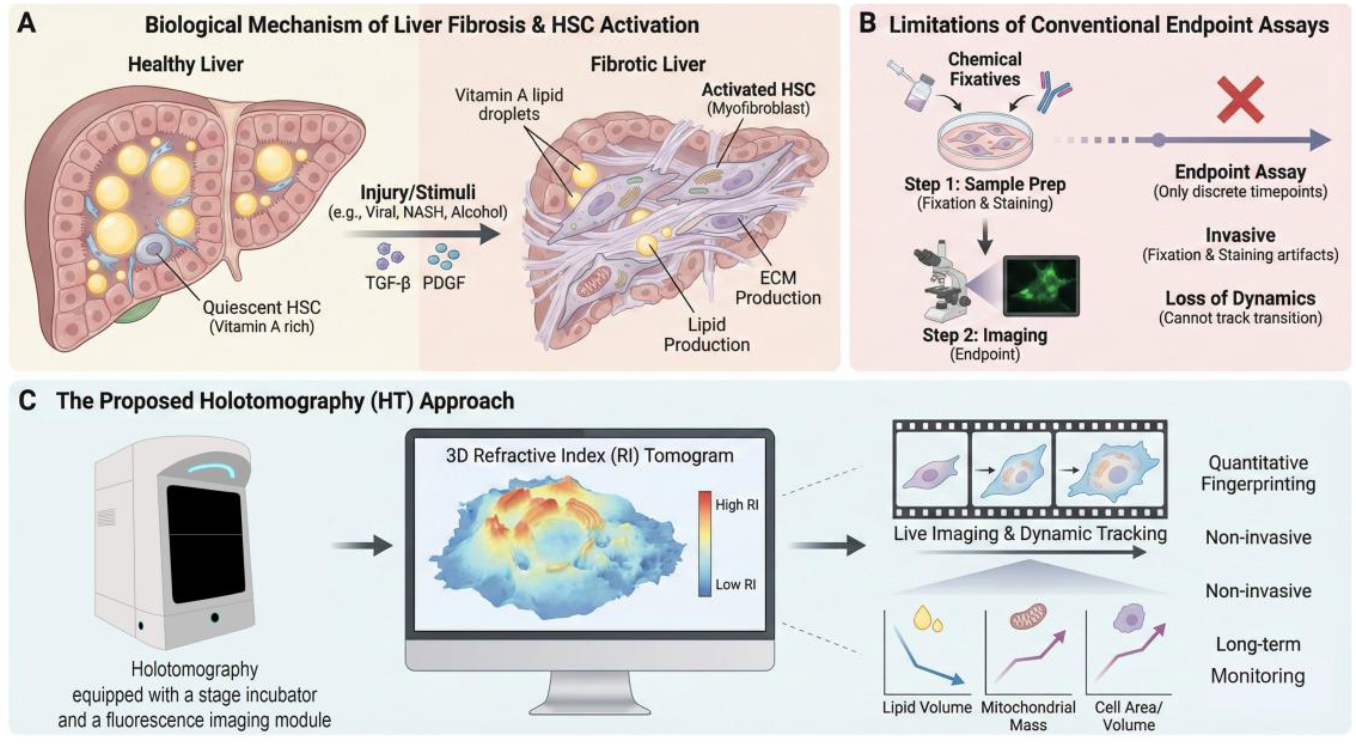
Conceptual framework for label-free, dynamic phenotypic profiling of hepatic stellate cell activation by holotomography. **(A)** Schematic illustration of hepatic stellate cell (HSC) activation during liver fibrosis. In healthy liver tissue, quiescent HSCs store vitamin A–rich lipid droplets. Upon chronic injury or profibrotic stimuli (e.g., TGF-β, PDGF), HSCs undergo activation into myofibroblast-like cells, characterized by lipid droplet depletion, cytoskeletal remodeling, and excessive extracellular matrix (ECM) production. **(B)** Limitations of conventional endpoint assays for studying HSC activation. Standard workflows rely on fixation and staining at discrete time points, providing only static snapshots and preventing continuous tracking of activation dynamics. These approaches are invasive, labor-intensive, and poorly suited for capturing gradual phenotypic transitions in living cells. **(C)** Overview of the proposed holotomography (HT)-based approach for label-free phenotypic profiling. Live HSCs are imaged longitudinally using low-power illumination to reconstruct three-dimensional refractive index (RI) tomograms. Quantitative analysis of intrinsic biophysical parameters—including lipid droplet volume, mitochondrial mass, and cell morphology—enables non-invasive, long-term monitoring and quantitative fingerprinting of HSC activation without exogenous labels.

Here, we introduce a label-free, quantitative framework for profiling HSC activation based on three-dimensional HT combined with automated, AI-assisted analysis (Fig. 1B). By longitudinally imaging living cells and extracting multi-parametric biophysical descriptors at both whole-cell and organelle-resolved levels, this approach enables continuous monitoring of activation trajectories without perturbation. Correlative fluorescence imaging is used to validate the biological relevance of HT-derived features, while automated analysis establishes scalability and objectivity.

Using this framework, we demonstrate that HSC activation can be characterized as a continuous phenotypic transition defined by coordinated changes in cellular morphology, refractive index distributions, and organelle organization. Together, these results establish holotomography-based quantitative phenotyping as a generalizable strategy for studying dynamic cell-state transitions, bridging the gap between live-cell imaging and high-content phenotypic analysis.

## 2. Methods

### 2.1. Cell preparation and Activation protocol

Human hepatic stellate cells (LX-2) were cultured in Dulbecco’s Modified Eagle Medium (DMEM; Gibco) supplemented with 2% fetal bovine serum (FBS), 1% penicillin-streptomycin, and 2 mM L-glutamine under standard incubator conditions (37°C, 5% CO2). Cells were seeded onto glass-bottom culture dishes and allowed to adhere for 24 h prior to experimental manipulation.

To induce defined activation states, cells were assigned to one of three experimental conditions. Control (CTL) cells were maintained in standard culture medium. Serum-starved (STV) cells were incubated overnight in medium containing 0.1% FBS to establish a quiescent baseline. Activated (ACT) cells were serum-starved and subsequently treated with recombinant human TGF-β1 (2.5 ng mL^−1^) for 24 h to induce myofibroblast-like activation.

### 2.2. Holotomography image acquisition

Label-free three-dimensional RI imaging was performed using a HT microscope (HT-X1 Plus, Tomocube Inc.) equipped with an epifluorescence imaging module (FLX 100, Tomocube Inc.). The system reconstructs volumetric RI distributions from multiple measurements with various illumination patterns under low-power illumination, enabling non-invasive imaging of live cells [79–82]. The lateral and axial resolution of the HT system is 160 nm and 1 μm [83].

For longitudinal experiments, cells were imaged within an on-stage environmental chamber maintaining physiological temperature (37°C) and CO2 concentration (5%). Time-lapse 3D datasets were acquired every 15 min over a total duration of up to 42 h using a 40× objective lens (numerical aperture 0.95), covering a field of view of 308 µm × 308 µm. Imaging parameters were kept constant across all experimental conditions to ensure quantitative comparability.

### 2.3. Time-lapse experimental design

To resolve activation dynamics, imaging was initiated with an 18 h serum starvation period to establish a stable morphological and RI baseline. TGF-β1 was then added directly to the imaging dish without interrupting acquisition, and imaging was continued for an additional 24 h. This design enabled continuous tracking of the same cells throughout the activation process, allowing reconstruction of single-cell activation trajectories in three dimensions.

### 2.4. AI-assisted segmentation and quantitative analysis

Three-dimensional RI tomograms were analyzed using an AI-assisted analysis pipeline (TomoAnalysis™, version 2.2). Automated segmentation was applied to identify whole-cell volumes and major subcellular organelles, including mitochondria and lipid droplets, directly from intrinsic RI contrast.

From the segmented volumes, a comprehensive set of quantitative descriptors was extracted at both the whole-cell and organelle levels. Whole-cell metrics included projected area, volume, sphericity, mean RI, dry mass, and intracellular concentration. Organelle-resolved metrics included projected area, surface area, and volume for mitochondria and lipid droplets. All measurements were performed in three dimensions unless otherwise specified.

To ensure robustness, identical analysis pipelines and parameter settings were applied across all experimental groups without manual intervention or post hoc threshold adjustment.

### 2.4. Correlative immunofluorescence imaging

For molecular validation, cells were fixed following holotomography imaging and subjected to immunofluorescence (IF) staining. Cells were incubated with primary antibodies against α-smooth muscle actin (α-SMA) and Vimentin, followed by Alexa Fluor–conjugated secondary antibodies. Nuclei were counterstained with DAPI.

Fluorescence imaging was performed using an integrated fluorescence module (FLX 100) on the same microscope platform to enable spatial correspondence between HT and fluorescence datasets. Z-stacks were acquired with matched axial coverage to the HT volumes to facilitate direct correlative analysis.

### 2.5. Statistical analysis

Quantitative data were analyzed using GraphPad Prism (version 5.0). For comparisons between experimental groups, unpaired two-tailed Student’s t-tests were applied unless otherwise indicated. Data are reported as mean ± s.e.m. Statistical significance was defined as P < 0.05. Each condition represents analysis of more than 3,000 cells across three biological replicates and two independent experimental batches.

## 3. Results

### 3.1 Label-free visualization and molecular validation of hepatic stellate cell activation

To evaluate whether HT captures biologically meaningful features of HS) activation, we performed correlative imaging across control, serum-starved conditions, and TGF-β1– activated LX-2 cells (Fig. 2; see Methods). Three-dimensional R) tomograms provided label-free visualization of cellular and subcellular architecture, which was directly compared with IF staining of established activation markers.

**Figure 2.**
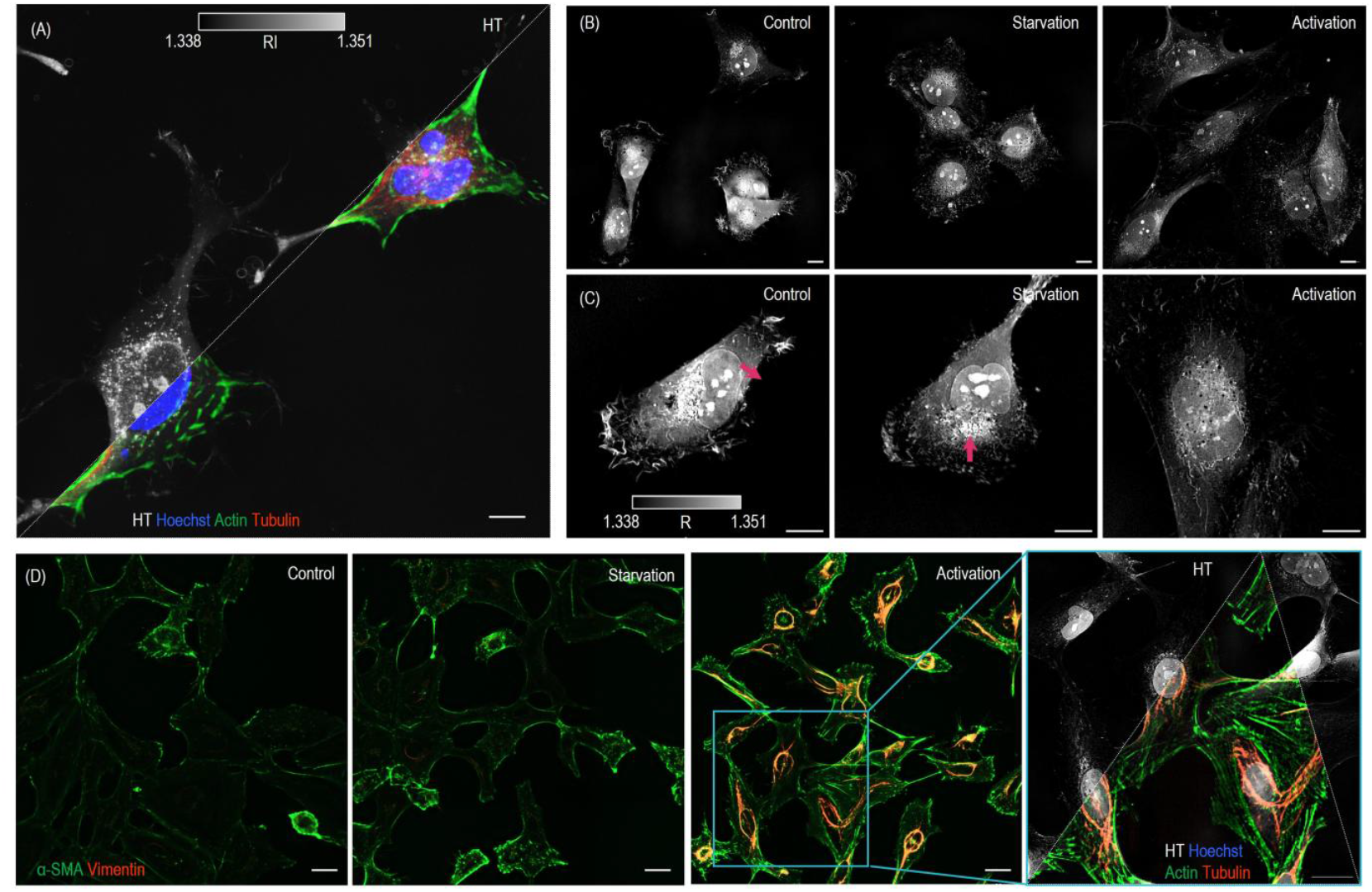
Label-free visualization and molecular validation of hepatic stellate cell activation using holotomography. **(A)** Correlative imaging workflow combining three-dimensional HT and confocal fluorescence microscopy. HT provides volumetric RI maps of live cells, which are subsequently compared with IF staining of established activation markers. **(B)** Representative XY slice views of LX-2 hepatic stellate cells under Control (CTL), Serum-Starved (STV), and TGF-β1-Activated (ACT) conditions. HT reveals pronounced morphological differences in the ACT group, including increased cell size and reduced overall intracellular RI, consistent with activation-associated cellular remodeling. **(C)** Magnified single-cell views highlighting subcellular refractive index features. Quiescent cells (CTL and STV) exhibit abundant high-RI granules (arrows), corresponding to vitamin A–rich lipid droplets, whereas these structures are markedly depleted in activated cells. The color bar indicates RI values. Scale bars, 10 μm. **(D)** Validation of HT-derived structural features by immunofluorescence. Maximum-intensity projection images show α-SMA (green) and Vimentin (red) expression across experimental groups, with strong induction and stress-fiber organization in the ACT condition. Correlative single-cell overlays demonstrate spatial alignment between filamentous structures observed in label-free HT images and fluorescently labeled cytoskeletal fibers, confirming that HT captures activation-associated cytoskeletal remodeling without exogenous labels. Scale bars, 35 μm (overview) and 25 μm (single-cell views).

Representative XY slices revealed clear activation-associated morphological differences in the TGF-β1–activated condition (Fig. 2B). Compared with control and serum-starved cells, activated HSCs exhibited pronounced cell spreading and increased projected area, accompanied by a reduction in overall intracellular RI. These changes are consistent with structural remodeling and altered intracellular composition during activation and were readily detectable without exogenous labels.

At the single-cell level, HT further resolved subcellular RI features characteristic of HSC state (Fig. 2C). Quiescent cells in the control and serum-starved groups displayed abundant high-RI granules, corresponding to vitamin A–rich lipid droplets that define the quiescent phenotype *[77]*. In contrast, these high-RI structures were markedly depleted in ACT cells, reflecting a canonical hallmark of HSC activation *[84]*. Importantly, these features were identified directly from intrinsic RI contrast, without prior knowledge of molecular markers *[85–88]*.

To validate that HT-derived structural features correspond to molecular activation, we performed correlative IF imaging of α-smooth muscle actin (α-SMA) and Vimentin (Fig. 2D). Consistent with established activation profiles, strong expression of both markers and organized stress-fiber architectures were observed predominantly in TGF-β1– activated cells, whereas control and serum-starved cells remained largely negative *[89]*. High-resolution correlative overlays demonstrated close spatial alignment between filamentous structures detected in label-free HT images and fluorescently labeled cytoskeletal fibers, confirming that HT faithfully captures activation-associated cytoskeletal remodeling.

Together, these results demonstrate that HT provides label-free access to both morphological and subcellular hallmarks of HSC activation, while maintaining direct correspondence with conventional molecular markers. This correlative validation establishes HT as a reliable, non-invasive modality for visualizing activation-related structural remodeling in living cells, forming the basis for subsequent longitudinal and quantitative analyses.

### 3.2 Real-time, label-free tracking of the hepatic stellate cell activation trajectory

While endpoint analyses confirmed distinct phenotypic differences between quiescent and activated HSCs, they do not capture the dynamic trajectory of activation. To directly observe this transition in living cells, we performed long-term time-lapse HT imaging to monitor HSC activation continuously over time (Fig. 3).

**Figure 3.**
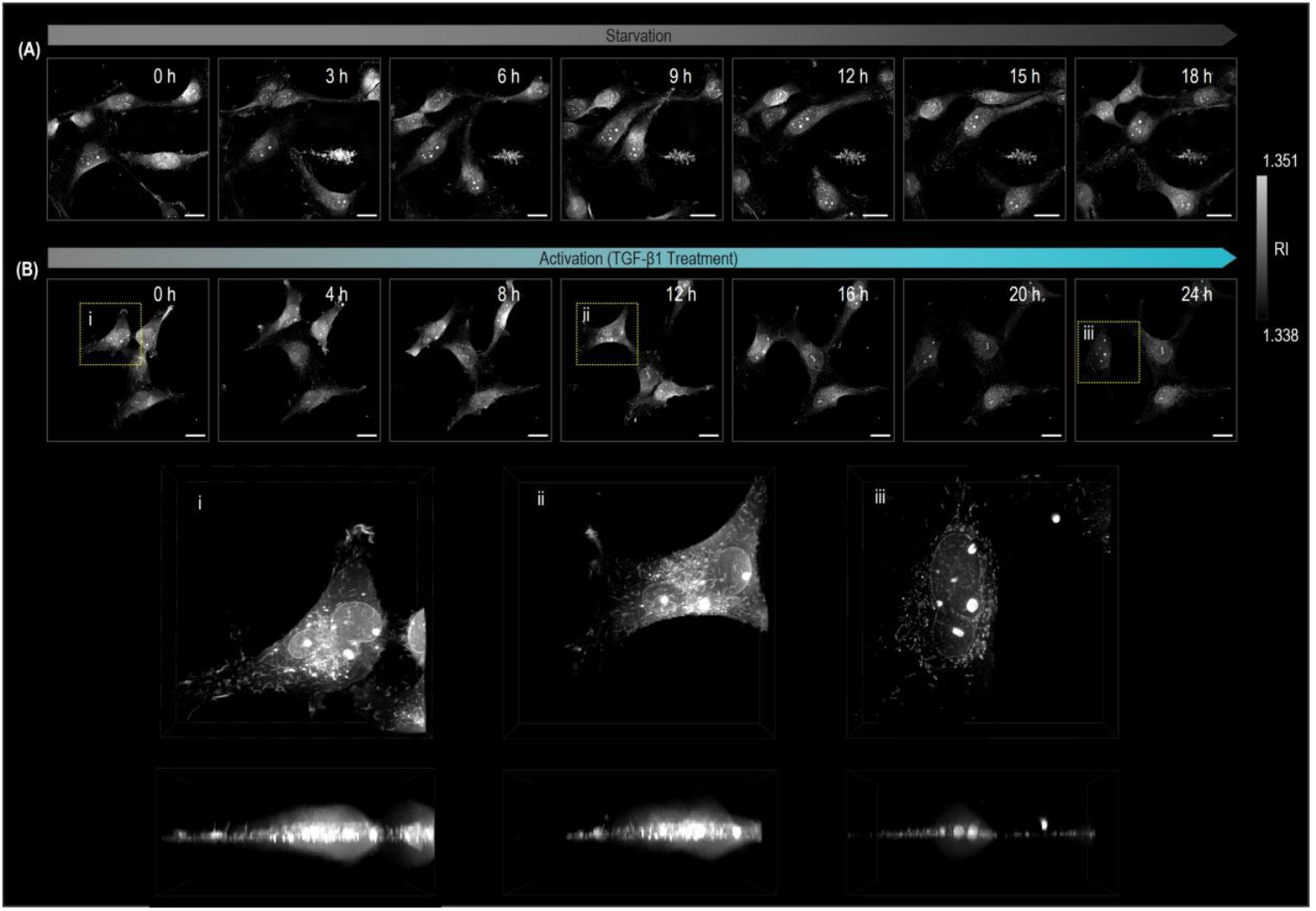
Real-time, label-free tracking of the hepatic stellate cell activation trajectory by HT. **(A)** Baseline characterization of quiescent hepatic stellate cells. Representative XY-slice time-lapse images acquired during an 18-hour serum starvation period establish a stable morphological and RI baseline prior to activation. Scale bars, 20 μm. **(B)** Longitudinal monitoring of activation kinetics following TGF-β1 stimulation. Time-lapse HT imaging over 24 hours reveals a gradual population-level transition characterized by progressive cell spreading and intracellular remodeling. The boxed region denotes a representative cell selected for detailed single-cell trajectory analysis in **(C)**. Scale bars, 20 μm. **(C)** Single-cell three-dimensional trajectory analysis. Volumetric RI renderings of the tracked cell at 0, 12, and 24 hours post-stimulation directly visualize activation-associated structural remodeling, including pronounced cell flattening and a marked reduction in cell height. These changes are difficult to capture using conventional two-dimensional endpoint assays.

Following an 18-hour serum starvation period, which established a stable morphological and RI baseline, cells exhibited minimal structural fluctuations at both the population and single-cell levels (Fig. 3A). This baseline provided a reference state for subsequent activation dynamics. Upon stimulation with TGF-β1, longitudinal HT imaging over the next 24 hours revealed a gradual, coordinated phenotypic transition across the cell population (Fig. 3B). Rather than an abrupt switch, activation unfolded as a continuous process characterized by progressive cell spreading and intracellular remodeling.

To resolve these dynamics at single-cell resolution, we tracked individual cells throughout the activation period and reconstructed their three-dimensional RI distributions at defined time points (Fig. 3C). Volumetric renderings directly visualized activation-associated structural remodeling, including pronounced cell flattening and a substantial reduction in cell height over time. These three-dimensional changes are difficult to infer from conventional two-dimensional microscopy or static endpoint assays, which cannot capture axial remodeling or continuous morphological trajectories.

Importantly, HT enabled these measurements without exogenous labels or fixation, allowing uninterrupted observation of the same living cells throughout the activation process. Together, these results demonstrate that holotomography provides real-time, label-free access to the dynamic trajectory of HSC activation, enabling direct visualization of gradual phenotypic transitions that are inaccessible to conventional endpoint-based approaches.

### 3.3 Automated, AI-assisted quantitative phenotypic fingerprinting of hepatic stellate cell activation

To translate label-free HT images into quantitative and scalable phenotypic descriptors, we applied an AI-assisted analysis pipeline implemented in TomoAnalysis™ to segment whole cells and subcellular organelles in three dimensions (Fig. 4A). This automated workflow enabled unbiased extraction of biophysical and organellar parameters directly from intrinsic RI tomograms, without manual annotation or fluorescent labeling [90].

**Figure 4.**
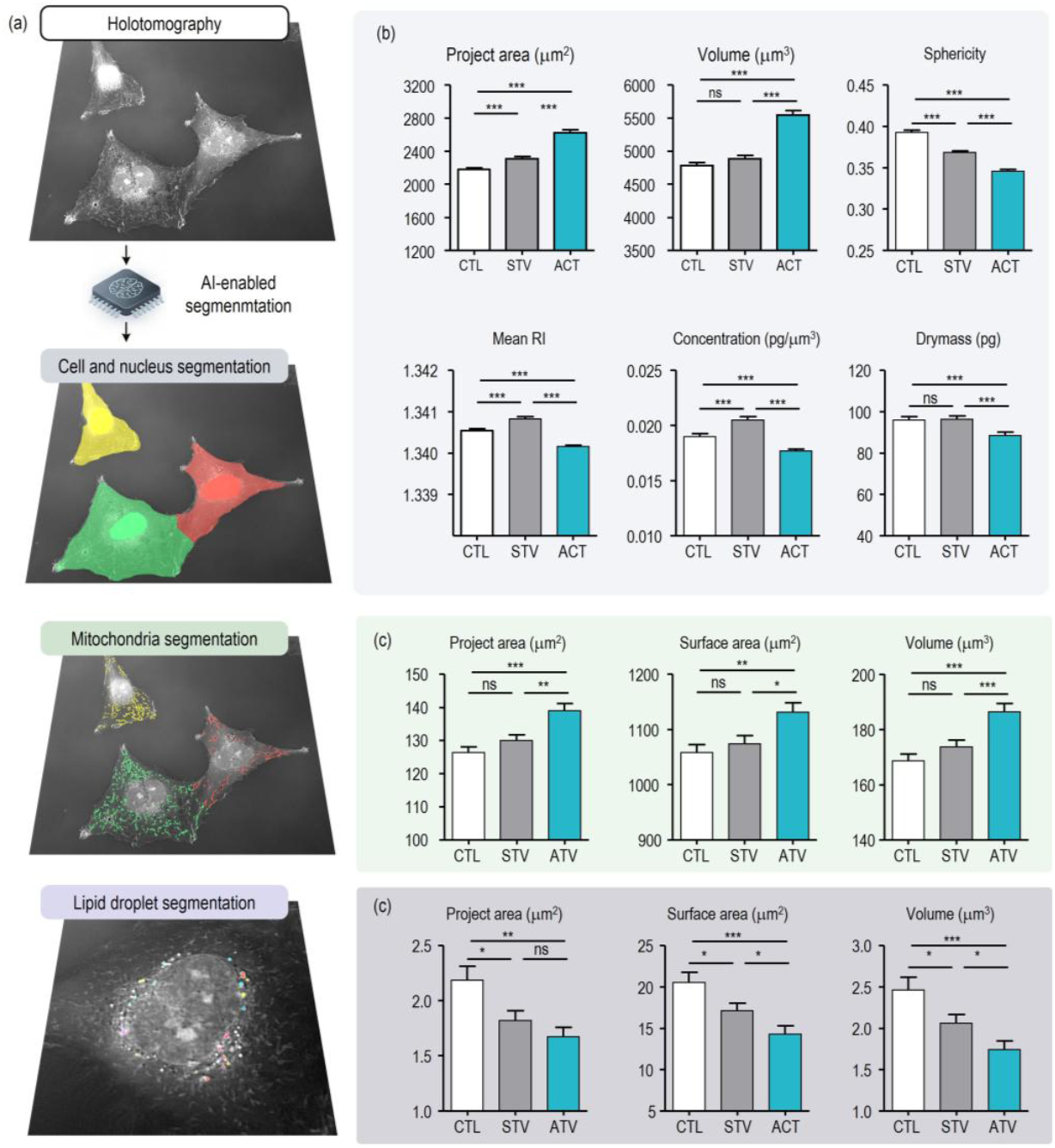
Automated, AI-assisted quantitative phenotypic fingerprinting of hepatic stellate cell activation from label-free holotomography. **(A)** AI-powered segmentation and analysis workflow implemented in TomoAnalysis™. Three-dimensional HT datasets are processed to independently segment whole-cell bodies, mitochondria, and lipid droplets, enabling systematic extraction of biophysical and organellar parameters from unlabeled cells. **(B)** Whole-cell–level biophysical profiling across Control (CTL), Serum-Starved (STV), and TGF-β1–Activated (ACT) conditions. Quantification of projected area, volume, sphericity, mean RI, dry mass, and intracellular concentration reveals coordinated shifts toward a hypertrophic and structurally remodeled phenotype in activated cells. **(C)** Organelle-resolved mitochondrial analysis. Per-cell measurements of mitochondrial projected area, surface area, and volume indicate activation-associated expansion of the mitochondrial population, consistent with altered metabolic demand during HSC activation. **(D)** Quantitative assessment of lipid droplet depletion. Per-cell lipid droplet projected area, surface area, and volume are significantly reduced in the ACT group, capturing a defining hallmark of HSC activation directly from intrinsic refractive index contrast. Data represent >3,000 cells per condition (three biological replicates across two independent batches). Statistical significance was assessed using unpaired two-tailed t-tests. Data are shown as mean ± s.e.m.; ns, not significant; *, *P* < 0.05; **, *P* < 0.01; ***, *P* < 0.001.

Whole-cell–level analysis revealed coordinated biophysical shifts associated with HSC activation (Fig. 4B). Compared with control and serum-starved conditions, TGF-β1–activated cells exhibited significant increases in projected area and volume, consistent with cellular hypertrophy. These morphological changes were accompanied by a reduction in mean intracellular RI, dry mass [91,92], and intracellular concentration, indicating altered intracellular composition during activation. A concurrent decrease in sphericity quantitatively captured the transition from compact to spread cell morphologies observed in label-free images.

Beyond whole-cell metrics, organelle-resolved analysis provided additional discriminatory power (Fig. 4C,D). Mitochondrial segmentation revealed significant increases in mitochondrial projected area, surface area, and volume in ACT cells, reflecting activation-associated expansion of the mitochondrial population. In contrast, lipid droplet analysis showed a pronounced reduction in lipid droplet abundance and volume, quantitatively capturing a defining hallmark of HSC activation directly from intrinsic RI contrast. Notably, these organelle-level measurements were derived without molecular labeling, relying solely on physical contrast.

Across all conditions, the combined set of whole-cell and organellar parameters defined a multi-dimensional biophysical fingerprint that robustly distinguished quiescent and activated HSC states. Analysis was performed on more than 3,000 cells per condition across independent biological replicates, demonstrating the scalability and robustness of the workflow. Together, these results establish that AI-assisted HT enables automated, high-throughput quantitative phenotypic profiling of cell-state transitions from label-free three-dimensional imaging data.

## 4. Discussions and Conclusion

In this study, we present a label-free, quantitative framework for profiling HSC activation based on three-dimensional HT. By combining intrinsic RI imaging with automated, AI-assisted analysis, this approach enables non-invasive visualization, longitudinal tracking, and multi-parametric quantification of cell-state transitions in living cells. Rather than relying on discrete, endpoint-based assays, HT provides continuous access to phenotypic dynamics across spatial and temporal scales.

A central advantage of this framework lies in its ability to capture biologically meaningful structural features without exogenous labels. Correlative imaging confirmed that HT-resolved morphological and subcellular features correspond closely to established molecular markers of HSC activation, including cytoskeletal remodeling and lipid droplet depletion. Importantly, these features are extracted directly from intrinsic physical contrast, avoiding perturbations associated with fixation, staining, or genetic modification. This establishes HT as a reliable modality for observing activation-associated remodeling while preserving native cellular states.

Beyond static observation, time-lapse HT enabled direct visualization of the activation trajectory at single-cell resolution. Activation was revealed as a gradual, continuous process characterized by coordinated changes in cell morphology and three-dimensional architecture, rather than an abrupt state switch. Such axial remodeling and continuous trajectories are difficult to infer from conventional two-dimensional microscopy or snapshot-based assays, highlighting the added value of volumetric, longitudinal imaging for studying dynamic cell-state transitions.

The integration of AI-assisted segmentation further extends HT from an imaging modality to a quantitative phenotyping platform. Automated extraction of whole-cell and organelle-resolved parameters yielded a multidimensional biophysical fingerprint that robustly distinguished quiescent and activated HSC states across thousands of cells. This scalability and objectivity enable systematic comparison of phenotypic states with minimal manual intervention, supporting applications that require high-content, high-throughput analysis.

While demonstrated here in the context of HSC activation, this framework is broadly applicable to other cell-state transitions involving coordinated morphological and metabolic remodeling, such as differentiation, immune activation, and drug-induced responses [93–97]. Because the approach is label-free and compatible with long-term live-cell imaging, it is particularly well suited for phenotypic screening workflows where perturbation-free monitoring and temporal resolution are critical.

Several limitations merit consideration. Although RI-based measurements capture integrated biophysical properties, they do not directly report molecular identity. As such, HT is complementary to, rather than a replacement for, molecular assays when specific biochemical pathways must be interrogated. In addition, interpretation of organelle-resolved features depends on robust segmentation performance, which may vary with imaging conditions or cell types. Continued refinement of analysis models and cross-validation with orthogonal modalities will further strengthen the generalizability of the approach.

Beyond the present demonstration, the proposed framework is inherently extensible across both analytical and biological dimensions. Integration with advanced AI-based analysis enables further expansion, for example by performing individual cell segmentation followed by extraction of intracellular biophysical parameters to support data-driven classification of cell states [98,99]. In parallel, virtual staining approaches could be leveraged to generate molecularly specific representations that can be analyzed in a correlative manner with quantitative holotomography-derived metrics, enabling joint interpretation of physical and inferred molecular features [100,101]. Although this study focuses on monolayer cell cultures, the methodology is directly compatible with more complex systems, including three-dimensional cultures, patient-derived organoids, and tissue specimens [43,102–106], where label-free, volumetric, and quantitative phenotyping is particularly advantageous. These extensions position the framework as a flexible foundation for multimodal, AI-enabled phenotypic analysis across diverse biological contexts.

In summary, this work establishes holotomography combined with automated analysis as a general, label-free strategy for quantitative phenotypic profiling of dynamic cell-state transitions. By enabling continuous, non-invasive access to three-dimensional cellular remodeling at scale, this framework bridges the gap between live-cell imaging and quantitative phenotyping, providing a foundation for standardized, high-content analysis in modern cell biology and translational research.

## Acknowledgments

We acknowledge Korea Basic Science Institute (Ochang Center) for valuable scientific discussions and expert advice that supported this application note.

## Conflicts of interest

All the authors have financial interests in Tomocube Inc., a company that commercializes HT system.

## Funding

This work was supported by the National Basic Science Research Program through the National Research Foundation of Korea (NRF) funded by the Ministry of Science and ICT (RS-2024-00442348), Korea Institute for Advancement of Technology (KIAT) through the International Cooperative R&D program (P0028463).

## References

1. C. Trapnell, D. Cacchiarelli, J. Grimsby, P. Pokharel, S. Li, M. Morse, N. J. Lennon, K. J. Livak, T. S. Mikkelsen, and J. L. Rinn, “The dynamics and regulators of cell fate decisions are revealed by pseudotemporal ordering of single cells,” Nat Biotechnol 32, 381–386 (2014).

2. L. Diamante and G. Martello, “Metabolic regulation in pluripotent stem cells,” Current Opinion in Genetics & Development 75, 101923 (2022).

3. N. Moris, C. Pina, and A. M. Arias, “Transition states and cell fate decisions in epigenetic landscapes,” Nat Rev Genet 17, 693–703 (2016).

4. S. J. Altschuler and L. F. Wu, “Cellular Heterogeneity: Do Differences Make a Difference?,” Cell 141, 559–563 (2010).

5. Y. Park, C. Depeursinge, and G. Popescu, “Quantitative phase imaging in biomedicine,” Nature Photon 12, 578–589 (2018).

6. M. Chen, H. Ma, X. Sun, M. Schwartz, R. E. Brand, J. Xu, D. S. Gotsis, P. Nguyen, B. A. Moore, L. Snyder, R. M. Brand, and Y. Liu, “Multimodal whole slide image processing pipeline for quantitative mapping of tissue architecture and tissue microenvironment,” npj Imaging 3, 26 (2025).

7. Y. Liu and S. Uttam, “Perspective on quantitative phase imaging to improve precision cancer medicine,” JBO 29, S22705 (2024).

8. G. Kim, H. Hugonnet, K. Kim, J.-H. Lee, S. S. Lee, J. Ha, C. Lee, H. Park, K.-J. Yoon, and Y. Shin, “Holotomography,” Nature Reviews Methods Primers 4, 51 (2024).

9. D. Pirone, C. Di Natale, M. Di Summa, N. Mosca, G. Giugliano, M. Schiavo, D. Florio, D. Marasco, P. L. Maffettone, L. Miccio, P. Memmolo, and P. Ferraro, “From genotype to phenotype: decoding mutations in blasts by holotomographic flow cytometry,” Light Sci Appl 14, 233 (2025).

10. D. Pirone, J. Lim, F. Merola, L. Miccio, M. Mugnano, V. Bianco, F. Cimmino, F. Visconte, A. Montella, M. Capasso, A. Iolascon, P. Memmolo, D. Psaltis, and P. Ferraro, “Stainfree identification of cell nuclei using tomographic phase microscopy in flow cytometry,” Nat. Photon. 16, 851–859 (2022).

11. S. Y. Lee, H. J. Park, C. Best-Popescu, S. Jang, and Y. K. Park, “The Effects of Ethanol on the Morphological and Biochemical Properties of Individual Human Red Blood Cells,” PLOS ONE 10, e0145327 (2015).

12. G. Kim, M. Lee, S. Youn, E. Lee, D. Kwon, J. Shin, S. Lee, Y. S. Lee, and Y. Park, “Measurements of three-dimensional refractive index tomography and membrane deformability of live erythrocytes from Pelophylax nigromaculatus,” Sci Rep 8, 9192 (2018).

13. H. Park, T. Ahn, K. Kim, S. Lee, S. Kook, D. Lee, I. B. Suh, S. Na, and Y. Park, “Three-dimensional refractive index tomograms and deformability of individual human red blood cells from cord blood of newborn infants and maternal blood,” J. Biomed. Opt 20, 111208 (2015).

14. M. Kim, W. S. Park, G. Kim, S. Oh, J. Do, J. Park, and Y. Park, “Label-free classification of cell death pathways via holotomography-based deep learning framework,” (n.d.).

15. S. Yoo, E. Yang, S.-H. Kim, H. Park, D. Kim, and M. L. Choi, “Organelle-Aware Representation Learning Enables Label-Free Detection of Mitochondrial Dysfunction in Live Human Neurons,” (n.d.).

16. J. Park, J. Lee, H.-J. Kim, S.-H. Chae, J. Shin, J.-H. Lee, Y. Cheon, Y. Jung, S.-K. Mun, J.-J. Kim, S.-H. Kim, G.-S. Hwang, and S. Lee, “Activation of lands cycle-mediated inflammation in living macrophages exposed to label-free particulate matter,” Journal of Hazardous Materials 499, 140027 (2025).

17. P. Anantha, A. Gupta, J. H. Kim, E. Saracino, P. Raj, I. Lucarini, S. Tanwar, J. Chen, L. Gu, J. Agrawal, A. Convertino, and I. Barman, “Disordered Glass Nanowire Substrates Produce in Vivo-Like Astrocyte Morphology Revealed by Low-Coherence Holotomography,” Advanced Science e13424 (2025).

18. J. Oh, H. Hugonnet, W. S. Park, and Y. Park, “Generalized reciprocal diffractive imaging for reference-free, single-shot quantitative phase microscopy,” Commun Phys 8, 383 (2025).

19. M. A. Ferrara, G. Preziosi, R. Boni, R. Ruggiero, and S. C. Gualandi, “Quantitative holographic analysis in stallion spermatozoa following cryopreservation,” Sci Rep 15, 43190 (2025).

20. J. Fu, Q. Ni, Y. Wu, A. Gupta, Z. Ge, H. Yang, Y. Afrida, I. Barman, and S. X. Sun, “Cells prioritize the regulation of cell mass density,” Science AdvAnceS (2025).

21. S. Kawagoe, M. Matsusaki, T. Mabuchi, Y. Ogasawara, K. Watanabe, K. Ishimori, and T. Saio, “Mechanistic Insights Into Oxidative Response of Heat Shock Factor 1 Condensates,” JACS Au 5, 606–617 (2025).

22. Y. Honda, D. Tokura, S. El Muttaqien, K. Konarita, Y. Kawashima, N. Nishiyama, and T. Nomoto, “Phenylboronic acid-based polymers exerting intracellular hydrophilic-to-hydrophobic conversion to retain within target cells,” Chemical Engineering Journal 527, 171908 (2026).

23. Y. Li, Y. Zhang, M. Chen, S. Antoku, J. Ding, K. Huang, G. G. Gundersen, and W. Chang, “Elevated SUN1 promotes migratory cell polarity defects through mechanically coupling microtubules to the nuclear lamina,” Commun Biol (2025).

24. K. Yu, S. T. Chua, A. Smith, A. G. Smith, T. Ellis, and S. Vignolini, “Cultivating Future Materials: Artificial Symbiosis for Bulk Production of Bacterial Cellulose Composites,” 2025.04.23.650277 (2025).

25. A. Tsukamura, H. Ariyama, N. Hayashi, S. Miyatake, S. Okado, S. Sultana, I. Terakado, T. Yamamoto, S. Yamanaka, S. Fujii, H. Hamanoue, R. Asano, T. Mizushima, N. Matsumoto, Y. Maruo, and M. Mori, “KNTC1 introduces segmental heterogeneity to mitochondria,” Disease Models & Mechanisms 18, DMM052063 (2025).

26. Y. Lee, W. H. Jung, K. Jeon, E. B. Choi, T. Ryu, C. Lee, D.-N. Kim, and D. J. Ahn, “Membrane-targeted DNA frameworks with biodegradability recover cellular function and morphology from frozen cells,” Trends in Biotechnology 43, 3196–3216 (2025).

27. P. Anantha, X. Wu, S. Elsaid, P. Raj, I. Barman, and S. S. Tee, “Sweet science: Exploring the impact of fructose and glucose on brown adipocyte differentiation using optical diffraction tomography,” (2024).

28. P. Rawat, T. Quaderer, I. Karemaker, S. S. Lee, F. Ulliana, Z. Kontarakis, J. E. Corn, and M. Peter, “1 Disruption of nucleolar integrity triggers cellular 2 quiescence through organelle rewiring and secretion,” (n.d.).

29. M. Machida, S. Kajimoto, R. Shibuya, M. Isono, M. Watabe, Y. Oma, K. Hibino, K. Fujii, M. Okumura, M. Harata, A. Shibata, and T. Nakabayashi, “Lipids Are Involved in Heterochromatin Condensation: A Quantitative Raman and Brillouin Microscopy Study,” (2025).

30. S. Kroschwald, F. Uliana, C. Wilson-Zbinden, A. Timofiiva, J. Zhou, S. S. Lee, L. Gillet, M. Zanella, A. Othman, R. Mezzenga, and M. Peter, “PKA regulates stress granule maturation to allow timely recovery after prolonged starvation,” 2025.07.06.663161 (2025).

31. J. Cho, M. J. Lee, J. Park, J. Lee, S. Lee, C. Chung, B.-K. Koo, Y. Park, and J. Cho, “Label-free, High-Resolution 3D Imaging and Machine Learning Analysis of Intestinal Organoids via Low-Coherence Holotomography,” Journal of Visualized Experiments (JoVE) e68529 (2025).

32. M.-T. Hong, G. Lee, and Y.-T. Chang, “A Non-Invasive, Label-Free Method for Examining Tardigrade Anatomy Using Holotomography,” Tomography 11, 34 (2025).

33. K. Han, J. Choi, C. Kim, S. Kang, H. An, C.-G. Pack, J.-H. Ahn, H. Kwon, C. W. Kim, J. S. Song, T. W. Kim, E. Tak, and J. E. Kim, “Gelatin-Based Soft-Tissue Sarcoma Organoids Recapitulate Patient Tumor Characteristics,” Biomater Res 8, 0293 (2025).

34. M. Amirola-Martinez, T. Combriat, K. Ferencevic, I. Wilhelmsen, A. Dalmao-Fernandez, P. A. Olsen, J. Stokowiec, A. Aizenshtadt, and S. Krauss, “Aspects of zone-like identity and holotomographic tracking of human stem cell-derived liver sinusoidal endothelial cells,” Front. Cell Dev. Biol. 13, 1528991 (2025).

35. J. Hong, H. Hugonnet, C. M. Oh, W. S. Park, C. Lee, C. Lee, Y. W. Kim, W. Heo, S.-M. Hong, and Y. Park, “High-speed holotomography of live cells and tissues using multi-pattern sparse axial scanning,” Opt. Express 33, 45708 (2025).

36. H. Kim, S.-Y. Heo, Y.-I. Kim, D. Park, Monford Paul Abishek N, S. Hwang, Y. Lee, H. Jang, J.-W. Ahn, J. Ha, S. Park, H. Y. Ji, S. Kim, I. Choi, W. Kwon, J. Kim, K. Kim, J. Gil, B. Jeong, J. C. D. Lazarte, R. Rollon, J. H. Choi, E. H. Kim, S.-G. Jang, H. K. Kim, B.-Y. Jeon, G. Kayali, R. J. Webby, B.-K. Koo, and Y. K. Choi, “Diverse bat organoids provide pathophysiological models for zoonotic viruses,” Science 388, 756–762 (2025).

37. H. Miki, M. M. Gomez, A. Itani, D. Yamanaka, Y. Sato, A. Di Pietro, and N. Takeshita, “Cell wall remodeling in a fungal pathogen is required for hyphal growth into microspaces,” mBio 16, e01184–25 (2025).

38. J. Simińska-Stanny, P. Tournier, A. Shavandi, and S. J. Habib, “Geometrical Designs in Volumetric Bioprinting to Study Cellular Behaviors in Engineered Constructs,” Adv Healthcare Materials e03550 (2025).

39. E. Ouni, A. Peaucelle, R. Ghasemi, F. Facchinetti, M. Opitz, L. Bigot, A. Sauvat, O. Kepp, F. Jaulin, Y. Loriot, and K. Schauer, “Mechanosensitive interactions of tumoroids with an engineered environment promote cell proliferation and enhance drug response detection,” Cell Biomaterials 1, 100149 (2025).

40. A. E. Melik-Pashaev, D. K. Matveeva, S. V. Buravkov, D. A. Atyakshin, E. S. Kochetova, and E. R. Andreeva, “Microscopy and Image Analysis of ?ell-Derived Decellularized Extracellular Matrix,” Cell Tiss. Biol. 19, 33–47 (2025).

41. S. Jeon, S.-H. Jeong, M. H. Lee, J. W. Seo, D.-S. Kim, N. J. Bassous, J. A. Lozano Soto, C. Choi, M. L. Gonzalez, E. B. Nolasco Díaz, H. Kim, S. R. Shin, and J.-U. Park, “Sustained oxygen-releasing hydrogel implants enhance flap regeneration by promoting mitochondrial biogenesis under mild hypoxia,” Bioactive Materials 51, 559–574 (2025).

42. S. Park, S. Y. Sun, J. G. Son, S. Y. Lee, H. K. Shon, O. Kwon, M. Lee, K. Choi, J. Yeun, S. H. Yoon, M. Kim, M. Son, T. G. Lee, and S. G. Im, “Tailored Xenogeneic-Free Polymer Surface Promotes Dynamic Migration of Intestinal Stem Cells,” Advanced Materials e13371 (2025).

43. C. Oh, J. Cho, J. Park, H. Lee, and Y. Park, “Morphology-Preserving Holotomography: Quantitative Analysis of 3D Organoid Dynamics,” (2025).

44. Y. Oyama, M. Ohama, N. Yamada, N. Suzuki, K. Kimura, and Y. Hara, “Tetraploid Caenorhabditis elegans embryos exhibit enhanced tolerance to osmotic stress,” 2025.10.13.678412 (2025).

45. A. Lauriola, J.H. Enriqué Steinberg M. Sarubo, E. Maspero, F. A. Rossi, Y. Mouri, M. Pedretti, M. Hajisadeghian, V. Taibi, A. Vettori, N. Vitulo, M. Assfalg, M. D’Onofrio, M. Rossi, A. Yasue, A. Astegno, S. Polo, S. Santi, Y. Kudo, and D. Guardavaccaro, “The E3 ligase RNF32 controls the IκB kinase complex and NF-κB signaling in intestinal stem cells,” Molecular Cell 85, 4254-4267.e9 (2025).

46. S. W. Park, H. C. Moon, S. J. Hong, A. Choi, S.-L. Lee, D. H. Park, E. Shin, J. H. Jo, D. H. Koh, J. Lee, J.-U. Hou, and K. J. Lee, “Enhancing biliary tract cancer diagnosis using AI-driven 3D optical diffraction tomography,” Methods 241, 196–203 (2025).

47. J. Park, S.-J. Shin, G. Kim, H. Cho, D. Ryu, D. Ahn, J. E. Heo, J. R. Clemenceau, I. Barnfather, and M. Kim, “Revealing 3D microanatomical structures of unlabeled thick cancer tissues using holotomography and virtual H&E staining,” Nature communications 16, 4781 (2025).

48. P. L. Wang, N. A. Lester, E. N. Perrault, J. Su, D. Gong, C. Shiau, J. Cao, P. T. T. Nguyen, J. W. Bae, D. Olgun, H. I. Hoffman, A. Lam, J. Huang-Gao, S. Rahaman, J. A. Guo, J. L. Barth, N. Caldwell, P. Divakar, J. W. Reeves, A. Bahrami, S. He, M. Patrick, E. Miller, M. Ganci, G. C. Jaramillo, T. S. Hong, J. Y. Wo, H. Roberts, R. Weissleder, H. Choi, C. F. Castillo, K. Cormier, D. T. Ting, T. Jacks, L. Zheng, M. Hemberg, M. Mino-Kenudson, and W. L. Hwang, “The Pdgfd-Pdgfrb axis orchestrates tumor-nerve crosstalk in pancreatic cancer,” 2025.08.26.672505 (2025).

49. M. Cangkrama, H. Liu, X. Wu, J. Yates, J. Whipman, C. G. Gäbelein, M. Matsushita, L. Ferrarese, S. Sander, F. Castro-Giner, S. Asawa, M. K. Sznurkowska, M. Kopf, J. Dengjel, V. Boeva, N. Aceto, J. A. Vorholt, and S. Werner, “MIRO2-mediated mitochondrial transfer from cancer cells induces cancer-associated fibroblast differentiation,” Nat Cancer 6, 1714–1733 (2025).

50. L. Bianchi, A. Bresci, K. J. Kobayashi-Kirschvink, G. Paroni, P. Saccomandi, P. T. C. So, and J. W. Kang, “A Multimodal Imaging Framework to Advance Phenotyping of Living Label-free Breast Cancer Cells,” JoVE 68498 (2025).

51. H. Khadem, M. Mangini, M. A. Ferrara, A. C. De Luca, and G. Coppola, “Polarization-Sensitive Holotomography for Multidimensional Label-Free Imaging and Characterization of Lipid Droplets in Cancer Cells,” Advanced Science 12, e09420 (2025).

52. D. H. Min, D. Kim, S. T. Hong, J. Kim, M. J. Kim, S. Kwon, A. Kim, and J.-Y. Lee, “Bafilomycin A1 induces colon cancer cell death through impairment of the endolysosome system dependent on iron,” Sci Rep 15, 5148 (2025).

53. G. H. Baek, D. Kim, G. Son, H. Do, G.-B. Yeon, M. J. Lee, M. Ji, J.-H. Son, M. Ju, I. Ahn, C. S. Kang, H. Lee, S. Choi, J. M. Suh, J. Seo, F. H. Gage, M.-J. Paik, Y. Park, D.-S. Kim, and J. Han, “Differential effects of lithium on metabolic dysfunctions in astrocytes derived from bipolar disorder patients,” Mol Psychiatry 30, 5833–5848 (2025).

54. A. Rudawska, B. Szermer-Olearnik, A. Szczygieł, J. Mierzejewska, K. Węgierek-Ciura, P. Żeliszewska, D. Kozien, M. Chaszczewska-Markowska, Z. Adamczyk, P. Rusiniak, K. Wątor, A. Rapak, Z. Pędzich, and E. Pajtasz-Piasecka, “Functionalized Boron Carbide Nanoparticles as Active Boron Delivery Agents Dedicated to Boron Neutron Capture Therapy,” IJN Volume 20, 6637–6657 (2025).

55. T.-Y. Ha, Y. Kim, S. M. Lim, Y. Hong, and K.-A. Chang, “GPR40 agonist ameliorates neurodegeneration by regulating mitochondria dysfunction and NLRP3 inflammasome in Alzheimer’s disease animal models,” Biomedicine & Pharmacotherapy 192, 118678 (2025).

56. Y. Chen, R. Ballarò, M. Sans, F. I. Thege, M. Zuo, R. Dou, J. Min, M. Yip-Schneider, J. Zhang, R. Wu, E. Irajizad, Y. Makino, K. I. Rajapakshe, H. K. Rudsari, M. W. Hurd, R.A. León-Letelier, H. Katayama, E. Ostrin, J. Vykoukal, J. B. Dennison, K.-A. Do, S. M. Hanash, R. A. Wolff, P. A. Guerrero, M. Kim, C. M. Schmidt, A. Maitra, and J. F. Fahrmann, “Long-chain sulfatide enrichment is an actionable metabolic vulnerability in intraductal papillary mucinous neoplasm (IPMN)-associated pancreatic cancers,” Gut 74, 1638–1652 (2025).

57. A. Zhbanov, Y. S. Lee, M. Son, B. J. Kim, and S. Yang, “Microfluidic Electrochemical Impedance Sensor for Hematological Tests of Blood under Different Osmotic Conditions,” Anal. Chem. 97, 21249–21257 (2025).

58. K.-H. Chang, H.-C. Chen, C.-Y. Chen, S.-P. Tsai, M.-Y. Hsu, P.-Y. Wang, S.-Y. Wu, and C.-L. Su, “Natural lignan justicidin A-induced mitophagy as a targetable niche in bladder cancer,” Chemico-Biological Interactions 421, 111723 (2025).

59. J. Ngoenkam, D. Pejchang, T. Nuamchit, U. Wichai, S. Pongcharoen, T. Laorob, and P. Paensuwan, “Nitro Dihydrocapsaicin Attenuates Hyperosmotic Stress-Induced Inflammation in the Corneal Epithelial Cells via SIRT1/Nrf2/HO-1 Pathway,” Experimental Eye Research 261, 110680 (2025).

60. C. Kim, S. Hong, S. H. Ma, J. Lee, H. So, J. Y. Kim, E. Shin, K. Lee, S. Choi, J. Park, Y. Park, Y.-M. Kim, J. H. Kim, and J. Kim, “Replication stress–induced nuclear hypertrophy alters chromatin topology and impacts cancer cell fitness,” Proc. Natl. Acad. Sci. U.S.A. 122, e2424709122 (2025).

61. L. A. Osminkina, P. A. Tyurin-Kuzmin, M. V. Sumarokova, and A. A. Kudryavtsev, “The Impact of Silicon Nanoparticle Porosity on Their Ability to Sensitize Low-Intensity Medical Ultrasound,” Sovrem Tehnol Med 17, 40 (2025).

62. Z. Liu, C. Chu, Y. Chen, C. Chung, F. Mi, M. Ho, W. Hsu, M. Hsieh, M. Chiang, C. Huang, P. Shueng, C. Yang, C. Lee, and C. Lin, “YAP Expression Confers Therapeutic Vulnerability to Cuproptosis in Breast Cancer Cells by Regulating Copper Homeostasis,” Adv Healthcare Materials e02769 (2025).

63. M. Ko, J. Ha, S. Kwon, H. E. Lee, J. Y. Mun, D. Yoon, J. Yoo, H. Cho, M. Lee, Y. Lee, S. Bae, J. Y. Lee, J. Y. Kim, S. H. Shin, M. H. Moon, and H. J. Kwon, “Cryptotanshinone Targets HYOU1 to Rewire ER-Mitochondria Communication and Enhance Autophagy in Atherosclerosis,” Research Square (2025).

64. B. K. Choi, H. H. Yang, J. H. Kim, J. Hong, K. M. Kim, and Y. R. Park, “Deep-Learning Model for Central Nervous System Infection Diagnosis and Prognosis Using Label-Free 3D Immune-Cell Morphology in the Cerebrospinal Fluid,” Advanced Intelligent Systems 7, 2401145 (2025).

65. S. D. Kumar, J. Park, N. K. Radhakrishnan, Y. P. Aryal, G. Jeong, I. Pyo, B. Ganbaatar, C. W. Lee, S. Yang, Y. Shin, S. Subramaniyam, Y. Lim, S. Kim, S. Lee, S. Y. Shin, and S. Cho, “Novel Leech Antimicrobial Peptides, Hirunipins: Real-Time 3D Monitoring of Antimicrobial and Antibiofilm Mechanisms Using Optical Diffraction Tomography,” Advanced Science 12, 2409803 (2025).

66. P. J. Pietras, M. Chaszczewska-Markowska, D. Ghete, A. Tyczewska, and K. Bąkowska-Żywicka, “Saccharomyces cerevisiae recovery from various mild abiotic stresses: Viability, fitness, and high resolution three-dimensional morphology imaging,” Fungal Genetics and Biology 178, 103975 (2025).

67. H.-S. Park, H.-G. Choi, I.-T. Jang, T. A. Pham, Z. Jiang, Y.-J. Son, K. Kim, and H.-J. Kim, “Endogenous hepcidin plays an essential role in Mycobacterium tuberculosis Rv1876 antigen-induced antimicrobial activity in macrophages,” Emerging Microbes & Infections 14, 2539192 (2025).

68. M. A. Ferrara, E. Cavalletti, V. Bianco, L. Miccio, G. Coppola, P. Ferraro, and A. Sardo, “Holographic tomography of the diatom Skeletonema pseudocostatum used as a bioindicator of heavy metal-polluted waters,” PLoS One 20, e0322960 (2025).

69. W. Dellisanti*, S. Murthy*, E. Bollati, S. P. Sandberg, and M. Kühl, “Moderate levels of dissolved iron stimulate cellular growth and increase lipid storage in Symbiodinium sp.,” (2024).

70. J. Park, D. D. Kang, H. Kim, J. H. Oh, and Y. Park, “Peptide PN5 from Pinus densiflora Confers in Vivo Protection Against Multidrug-Resistant Salmonella Typhimurium Through Membrane Disruption,” ACS Omega acsomega.5c06012 (2025).

71. S. Hong, Y.-E. Jeon, H. Kim, J. Hong, D. Lee, J. H. Park, J. Y. Jung, S. Cha, P. Lee, and J.-S. Hahn, “Pressing extraction: A novel and environmentally sustainable approach to microbial lipid extraction from Yarrowia lipolytica,” Separation and Purification Technology 377, 134148 (2025).

72. F. Salvà-Serra, P. Nimje, B. Piñeiro-Iglesias, L. A. Alarcón, S. Cardew, E. Inganäs, S. Jensie-Markopoulos, M. Ohlén, H.-S. Sailer, C. Unosson, V. Fernández-Juárez, C. O. Pacherres, M. Kühl, E. R. B. Moore, and N. P. Marathe, “Description of Pseudomonas imrae sp. nov., carrying a novel class C β-lactamase gene variant, isolated from gut samples of Atlantic mackerel (Scomber scombrus),” Front. Microbiol. 16, 1530878 (2025).

73. J. C. Caicedo, S. Cooper, F. Heigwer, S. Warchal, P. Qiu, C. Molnar, A. S. Vasilevich, J. D. Barry, H. S. Bansal, O. Kraus, M. Wawer, L. Paavolainen, M. D. Herrmann, M. Rohban, J. Hung, H. Hennig, J. Concannon, I. Smith, P. A. Clemons, S. Singh, P. Rees, P. Horvath, R. G. Linington, and A. E. Carpenter, “Data-analysis strategies for image-based cell profiling,” Nat Methods 14, 849–863 (2017).

74. S. L. Friedman, “Hepatic Stellate Cells: Protean, Multifunctional, and Enigmatic Cells of the Liver,” Physiological Reviews 88, 125–172 (2008).

75. J.-T. Li, Z.-X. Liao, J. Ping, D. Xu, and H. Wang, “Molecular mechanism of hepatic stellate cell activation and antifibrotic therapeutic strategies,” J Gastroenterol 43, 419–428 (2008).

76. D. A. Mann and D. E. Smart, “Transcriptional regulation of hepatic stellate cell activation,” Gut 50, 891–896 (2002).

77. W. S. Blaner, S. M. O’Byrne, N. Wongsiriroj, J. Kluwe, D. M. D’Ambrosio, H. Jiang, R. F. Schwabe, E. M. C. Hillman, R. Piantedosi, and J. Libien, “Hepatic stellate cell lipid droplets: A specialized lipid droplet for retinoid storage,” Biochimica et Biophysica Acta (BBA) - Molecular and Cell Biology of Lipids 1791, 467–473 (2009).

78. T. Kisseleva and D. A. Brenner, “Hepatic stellate cells and the reversal of fibrosis,” Journal of Gastroenterology and Hepatology 21, S84–S87 (2006).

79. Y. Chung, H. Hugonnet, S.-M. Hong, and Y. Park, “Fourier space aberration correction for high resolution refractive index imaging using incoherent light,” Opt. Express, OE 32, 18790– 18799 (2024).

80. H. Hugonnet, M. J. Lee, and Y. K. Park, “Quantitative phase and refractive index imaging of 3D objects via optical transfer function reshaping,” Opt. Express 30, 13802–13809 (2022).

81. H. Hugonnet, M. Lee, and Y. Park, “Optimizing illumination in three-dimensional deconvolution microscopy for accurate refractive index tomography,” Opt. Express 29, 6293–6301 (2021).

82. Y. Baek and Y. Park, “Intensity-based holographic imaging via space-domain Kramers–Kronig relations,” Nat. Photonics 15, 354–360 (2021).

83. C. Park, S. Shin, and Y. Park, “Generalized quantification of three-dimensional resolution in optical diffraction tomography using the projection of maximal spatial bandwidths,” Journal of the Optical Society of America A 35, 1891–1898 (2018).

84. T. Tsuchida and S. L. Friedman, “Mechanisms of hepatic stellate cell activation,” Nat Rev Gastroenterol Hepatol 14, 397–411 (2017).

85. M. Schürmann, J. Scholze, P. Müller, J. Guck, and C. J. Chan, “Cell nuclei have lower refractive index and mass density than cytoplasm,” Journal of Biophotonics 9, 1068–1076 (2016).

86. K. Kim, S. Lee, J. Yoon, J. Heo, C. Choi, and Y. Park, “Three-dimensional label-free imaging and quantification of lipid droplets in live hepatocytes,” Sci Rep 6, 36815 (2016).

87. S. Park, J. W. Ahn, Y. Jo, H.-Y. Kang, H. J. Kim, Y. Cheon, J. W. Kim, Y. Park, S. Lee, and K. Park, “Label-Free Tomographic Imaging of Lipid Droplets in Foam Cells for Machine-Learning-Assisted Therapeutic Evaluation of Targeted Nanodrugs,” ACS Nano 14, 1856–1865 (2020).

88. H. Kim, S. Oh, S. Lee, K. suk Lee, and Y. Park, “Recent advances in label-free imaging and quantification techniques for the study of lipid droplets in cells,” Current Opinion in Cell Biology 87, 102342 (2024).

89. A. Desmoulière, I. A. Darby, and G. Gabbiani, “Normal and Pathologic Soft Tissue Remodeling: Role of the Myofibroblast, with Special Emphasis on Liver and Kidney Fibrosis,” Laboratory Investigation 83, 1689–1707 (2003).

90. J. Park, B. Bai, D. Ryu, T. Liu, C. Lee, Y. Luo, M. J. Lee, L. Huang, J. Shin, and Y. Zhang, “Artificial intelligence-enabled quantitative phase imaging methods for life sciences,” Nature methods 20, 1645–1660 (2023).

91. R. Barer, “Determination of dry mass, thickness, solid and water concentration in living cells,” Nature 172, 1097–1098 (1953).

92. G. Popescu, Y. Park, N. Lue, C. Best-Popescu, L. Deflores, R. R. Dasari, M. S. Feld, and K. Badizadegan, “Optical imaging of cell mass and growth dynamics,” American Journal of Physiology-Cell Physiology 295, C538–C544 (2008).

93. M. D. Buck, D. O’Sullivan, and E. L. Pearce, “T cell metabolism drives immunity,” J Exp Med 212, 1345–1360 (2015).

94. E. Y. Jeong, H.-J. Kim, S. Lee, Y. Park, and Y. M. Kim, “Label-free long-term measurements of adipocyte differentiation from patient-driven fibroblasts and quantitative analyses of in situ lipid droplet generation,” J. Opt. Soc. Am. A 41, C125–C136 (2024).

95. P. Anantha, Z. Liu, P. Raj, and I. Barman, “Optical diffraction tomography and Raman spectroscopy reveal distinct cellular phenotypes during white and brown adipocyte differentiation,” Biosensors and Bioelectronics 235, 115388 (2023).

96. M. Cangkrama, H. Liu, X. Wu, J. Yates, J. Whipman, C. G. Gäbelein, M. Matsushita, L. Ferrarese, S. Sander, F. Castro-Giner, S. Asawa, M. K. Sznurkowska, M. Kopf, J. Dengjel, V. Boeva, N. Aceto, J. A. Vorholt, and S. Werner, “MIRO2-mediated mitochondrial transfer from cancer cells induces cancer-associated fibroblast differentiation,” Nat Cancer (2025).

97. J. Choi, S. Kang, H.-I. An, C.-E. Kim, S. Lee, C.-G. Pack, Y.-I. Yoon, H. Jin, Y.-P. Cho, C. J. Kim, J.-M. Namgoong, J. K. Kim, and E. Tak, “Fasudil and viscosity of gelatin promote hepatic differentiation by regulating organelles in human umbilical cord matrix-mesenchymal stem cells,” Stem Cell Res Ther 15, 229 (2024).

98. M. Lee, Y.-H. Lee, J. Song, G. Kim, Y. Jo, H. Min, C. H. Kim, and Y. Park, “Deep-learning-based three-dimensional label-free tracking and analysis of immunological synapses of CAR-T cells,” Elife 9, e49023 (2020).

99. H. Kim, G. Kim, H. Park, M. J. Lee, Y. Park, and S. Jang, “Integrating holotomography and deep learning for rapid detection of NPM1 mutations in AML,” Scientific reports 14, 23780 (2024).

100. Y. Jo, H. Cho, W. S. Park, G. Kim, D. Ryu, Y. S. Kim, M. Lee, S. Park, M. J. Lee, H. Joo, H. Jo, S. Lee, S. Lee, H. Min, W. D. Heo, and Y. Park, “Label-free multiplexed microtomography of endogenous subcellular dynamics using generalizable deep learning,” Nat Cell Biol 23, 1329–1337 (2021).

101. J. Park, S.-J. Shin, G. Kim, H. Cho, D. Ryu, D. Ahn, J. E. Heo, J. R. Clemenceau, I. Barnfather, M. Kim, I. Jang, J.-Y. Sung, J. H. Park, H. Min, K. S. Lee, N. H. Cho, T. H. Hwang, and Y. Park, “Revealing 3D microanatomical structures of unlabeled thick cancer tissues using holotomography and virtual H&E staining,” Nat Commun 16, 4781 (2025).

102. M. J. Lee, J. Lee, J. Ha, G. Kim, H.-J. Kim, S. Lee, B.-K. Koo, and Y. Park, “Long-term three-dimensional high-resolution imaging of live unlabeled small intestinal organoids via low-coherence holotomography,” Exp Mol Med 56, 2162–2170 (2024).

103. J. Cho, M. J. Lee, J. Park, J. Lee, S. Lee, C. Chung, B.-K. Koo, and Y. Park, “Label-free, High-Resolution 3D Imaging and Machine Learning Analysis of Intestinal Organoids via Low-Coherence Holotomography,” Journal of Visualized Experiments (JoVE) e68529 (2025).

104. D. Park, D. Lee, Y. Kim, Y. Park, Y.-J. Lee, J. E. Lee, M.-K. Yeo, M.-W. Kang, Y. Chong, S. J. Han, J. Choi, J.-E. Park, Y. Koh, J. Lee, Y. Park, R. Kim, J. S. Lee, J. Choi, S.-H. Lee, B. Ku, D. H. Kang, and C. Chung, “Cryobiopsy: A Breakthrough Strategy for Clinical Utilization of Lung Cancer Organoids,” Cells 12, 1854 (2023).

105. A. J. Lee, H. Hugonnet, Y. S. Kim, J.-G. Kim, M. Lee, T. Ku, and Y. Park, “Volumetric Refractive Index Measurement and Quantitative Density Analysis of Mouse Brain Tissue with Sub-Micrometer Spatial Resolution,” Advanced Photonics Research 4, 2300112 (2023).

106. H. Hugonnet, Y. W. Kim, M. Lee, S. Shin, R. H. Hruban, S.-M. Hong, and Y. Park, “Multiscale label-free volumetric holographic histopathology of thick-tissue slides with subcellular resolution,” Adv. Photon. 3, 026004 (2021).

